# Comparative Molecular Docking and Toxicity Profiling of Buspirone and Tandospirone Targeting the HTR1A Receptor

**DOI:** 10.1101/2025.11.17.688977

**Authors:** Yuhan Lu

## Abstract

The human serotonin 1A (5-HT1A/HTR1A) receptor is a central target in the treatment of anxiety and mood disorders. However, ligand efficacy is sensitive to receptor conformation. Both Buspirone and Tandospirone are clinically relevant 5-HT1A agonists. This comparative in silico analysis was performed to examine their binding behavior and toxicity profiles. Molecular docking was performed on the serotonin-bound HTR1A receptor (PDB ID: 7E2Y) using SwissDock, and the toxicological predictions were generated with ProTox 3.0. Both ligands exhibited the greatest binding affinity for Chain A of the receptor (-6.14 kcal/mol for Buspirone and -5.88 kcal/mol for Tandospirone), suggesting a preferred receptor conformation that may mediate therapeutic efficacy. Buspirone demonstrated greater binding stability than Tandospirone across models. Both compounds demonstrated weak or unstable interactions for chains G and R. Predicted toxicity profiles revealed high probabilities of neurotoxicity and respiratory toxicity for both ligands, with blood–brain barrier penetration probabilities of 0.99 (Buspirone) and 1.00 (Tandospirone).

Additionally, Tandospirone showed potential immunotoxic effects (probability 0.73). These findings demonstrated that the receptor conformation in ligand binding enhances efficacy in drug design while reducing CNS-related adverse effects. Overall, this comparative study provides a starting point that may inform the design of next-generation serotonergic therapeutics for mood and anxiety disorders.

## 1 INTRODUCTION

The 5-hydroxytryptamine receptor 1A gene (*HTR1A* or *5-HTR1A*), or serotonin 1A receptor gene, is located on the reverse strand of chromosome 5 at position 5q12.3, spanning coordinates 63,957,874 to 63,962,445 (National Center for Biotechnology Information, 2024). The gene encodes the serotonin 1A receptor (HTR1A, 5-HTR1A, or 5HT1A receptor), a G protein-coupled receptor (GPCR) consisting of 422 amino acids, which is crucial for regulating serotonin signaling in the brain (2024). These receptors are found in key brain areas such as the amygdala, hippocampus, prefrontal cortex, and hypothalamus, which are essential for managing mood, emotions, and stress responses (Le François et al., 2008).

Mental health disorders are significantly influenced by serotonin. Only 2% of the serotonin found in the human body is found in the central nervous system (CNS), but this small fraction is essential for mental health and imbalances or defects can contribute to the development of mental health disorders such as addiction, depression, suicidal behaviors, panic disorders anxiety, schizophrenia, post-traumatic stress disorder, and extreme phobias (Lin et al., 2014). Polymorphisms in serotonin receptor genes, such as *5-HTR1A*, can alter the expression, function, or binding affinity of their protein products. These allelic variations can impact serotonin processing in the brain, potentially increasing susceptibility to mental health disorders (Takekita et al., 2016).

5-HT1A receptors work in two main ways: they function as presynaptic autoreceptors on neurons that release serotonin, modulating the amount of serotonin they release, and as postsynaptic receptors on other brain cells, influencing how they respond to serotonin signals. Impaired 5-HTR1A function has been observed in individuals with psychiatric disorders like major depressive disorder (MDD), anxiety, and schizophrenia (Drevets et al., 2007). This impairment usually involves increased density or activity of presynaptic 5-HT1A receptors, resulting in the inhibition of serotonin (5-HT) release, worsening of symptoms, and poor response to antidepressants (Celada et al., 2013). While presynaptic 5-HT1A receptors are associated with detrimental effects in mood disorders, postsynaptic 5-HT1A receptors are beneficial for the therapeutic effects of antidepressant drugs and the activation of these receptors can contribute to reduced symptoms of mood disorders (Celada et al., 2013). The dual roles of 5-HT1A receptors complicate the treatment of mood disorders, making it difficult to achieve the desired therapeutic effects without also encountering counterproductive outcomes (Celada et al., 2013).

Considering the significant role that the regulation of serotonin plays in mental health, studying serotonin receptors, including 5-HTR1A, is crucial. Understanding the structures of these receptors and their polymorphisms can lead to more effective treatments and interventions, potentially reducing the prevalence of mental health conditions and improving the quality of life for many individuals. Molecular docking studies are invaluable to this line of research. These studies can be used to simulate how potential drug candidates interact with the HTR1A receptor, helping to assess the binding affinity of compounds of interest and identify drug candidates to design targeted therapies that consider various receptor polymorphisms (Agu et al., 2023). This approach enhances the precision of treatments and aims to minimize side effects, contributing to more effective management of mental health disorders.

Studying drugs that act as agonists or antagonists specifically targeting either presynaptic or postsynaptic 5-HT1A receptors depending on the mood disorder and symptoms is crucial for developing more precise and effective treatments. In this study, we explore the potential of buspirone and tandospirone as candidates for modulating 5-HT1A receptor activity through molecular docking studies. Our goal is to assess how buspirone and tandospirone interact with 5 - HT1A receptors, their binding affinity, and their effects on serotonin signaling. We will also evaluate their toxicity to ensure safety.

## 2 LITERATURE REVIEW

The serotonin 1A receptor is significantly linked with the molecular pathology of anxiety, which has been reported in several species. Studies have shown in mice, that a complete knockdown of 5 - HT1A causes an increase in anxiety-driven behaviors. Whereas when the receptor is over-expressed, the fear driven reactions are reduced (Hettema et al., 2008). There is evidence that supports mood disorders, such as anxiety, stemming from a tryptophan deletion in 5-HT, which indicates that the dysregulation of 5-HT neurotransmission is a main defect (Garcia-garcia et al., 2013). Novel therapeutics, namely 5HT1A, aim to directly target the raphe nuclei somatodendritic autoreceptors or the cortical heteroreceptors.

Recent research has deepened our understanding of how variations in the HTR1A gene can influence the function of the 5-HT1A receptor, impacting an individual’s vulnerability to mood disorders. For instance, the C(-1019)G (rs6295) polymorphism has been found to alter how transcription factors bind to the gene, which in turn affects the expression of the receptor in both presynaptic and postsynaptic areas. This altered expression has been linked to anxiety and depression, with the G-allele being particularly associated with a higher risk of depression, panic disorder, and neuroticism (Le François et al., 2008).

Additionally, studies have shown that the interaction between HTR1A and other genetic variations, such as those in the serotonin transporter gene (5-HTT), can significantly influence how well patients respond to antidepressants. For example, certain combinations of HTR1A and 5-HTT polymorphisms have been linked to a better response to mirtazapine in people with major depressive disorder, suggesting that these genetic markers could be useful in tailoring personalized treatments (Chang et al., 2014).

In terms of new treatment options, drugs that specifically target the 5-HT1A receptor are showing promise for treating anxiety and mood disorders. The effectiveness of partial agonists like perospirone and aripiprazole, which target this receptor, has been found to be related to specific HTR1A polymorphisms, highlighting the receptor’s role in influencing treatment outcomes (Takekita et al., 2015). Furthermore, recent studies suggest that the methylation status of HTR1A and related genes, when combined with factors like stress and genetic background, can play a key role in how effective antidepressants are. It extends exciting possibilities for epigenetic-based treatments (Xu et al., 2022).

## 3 COMPOUND SELECTION JUSTIFICATION

### Buspirone

Buspirone is a drug that has been identified for the treatment of generalized anxiety disorders. The compound binds to 5-HT1A receptors on both presynaptic neurons in dorsal raphe and postsynaptic neurons in the hippocampus (National Center for Biotechnology Information, 2024). It is considered an agonist, meaning it will increase serotonin receptor activity.

### Tandospirone

Tandospirone is a multifunctional drug used to treat anxiety and depression and currently enrolled in clinical trials for the treatment of Schizophrenia. The compound is a dicarboximide and a potent, partial agonist to the 5-HT1A receptor (National Center for Biotechnology Information, 2024).

## 4 METHODS (DOCKING & TOXICITY PREDICTION)

### Docking

Molecular docking was performed on the protein serotonin-bound Serotonin 1A (5HT1A) receptor-Gi protein complex with the ligands Buspirone and Tandospirone independently. Each of the 4 chains in the protein were docked separately. The 3D structure of the protein of interest as well as the ligands were visualized with UCSF chimera version 1.18. Molecular docking data was collected from the SwissDock platform using Docking with AutoDock Vina (Bugnon et al 2024). The data was analyzed by the resulting calculated affinity (kcal/mol). The 3D structure of 5-HT1A (7E2Y) was pulled from RCSB Protein Data Bank. The structure of Buspirone (CHEBI:3223) and Tandospirone (CHEBI:145673) were obtained from the Chemical Entities of Biological Interest (ChEBI).

### Toxicity Prediction

Toxicity predictions for both Buspirone and Tandospirone were completed with the ProTox 3.0-Prediction of TOXicity of Chemicals. The Canonical SMILES were used from ChEBI and all models were selected under the categories Organ toxicity, toxicity endpoints, molecular initiating events, Tox21 Nuclear receptor signaling pathways, Tox21 Stress response pathways, and Metabolism. The predicted toxicity class, the LD50, as well as the probability of an active toxic classification were analyzed.

## 5 RESULTS

### Docking

The 5HT1A consists of 4 chains (Figure 3). Chain A is Guanine nucleotide-binding protein G(i) subunit alpha-1, chain B is Guanine nucleotide-binding protein G(I)/G(S)/G(T) subunit beta-1, chain G is Guanine nucleotide-binding protein G(I)/G(S)/G(O) subunit gamma-2, and lastly chain R is the soluble cytochrome b562, 5-hydroxytryptamine receptor 1A (RCSB PDB, 2024).

**Figure 1.**
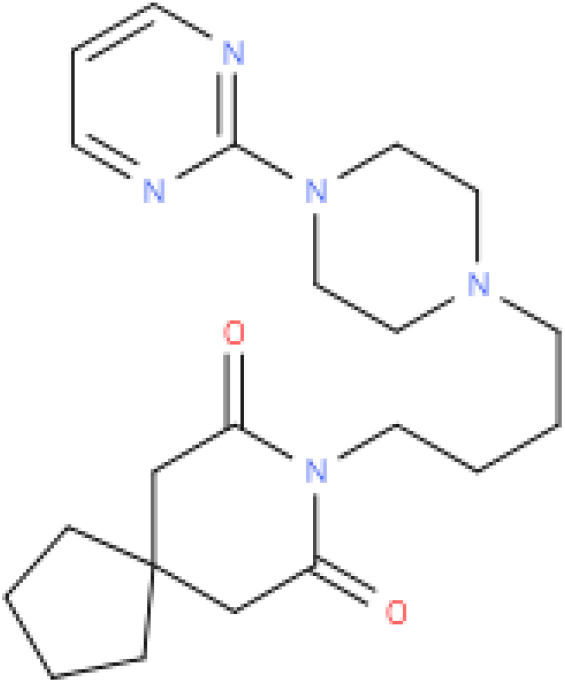
2D Buspirone structure. ChEBI:3223

**Figure 2.**
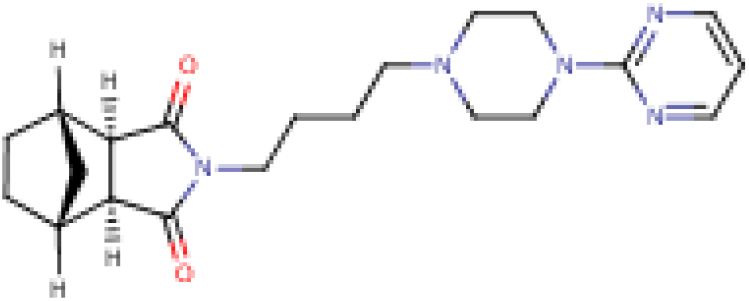
2D Tandospirone structure. ChEBI:145673

**Figure 3.**
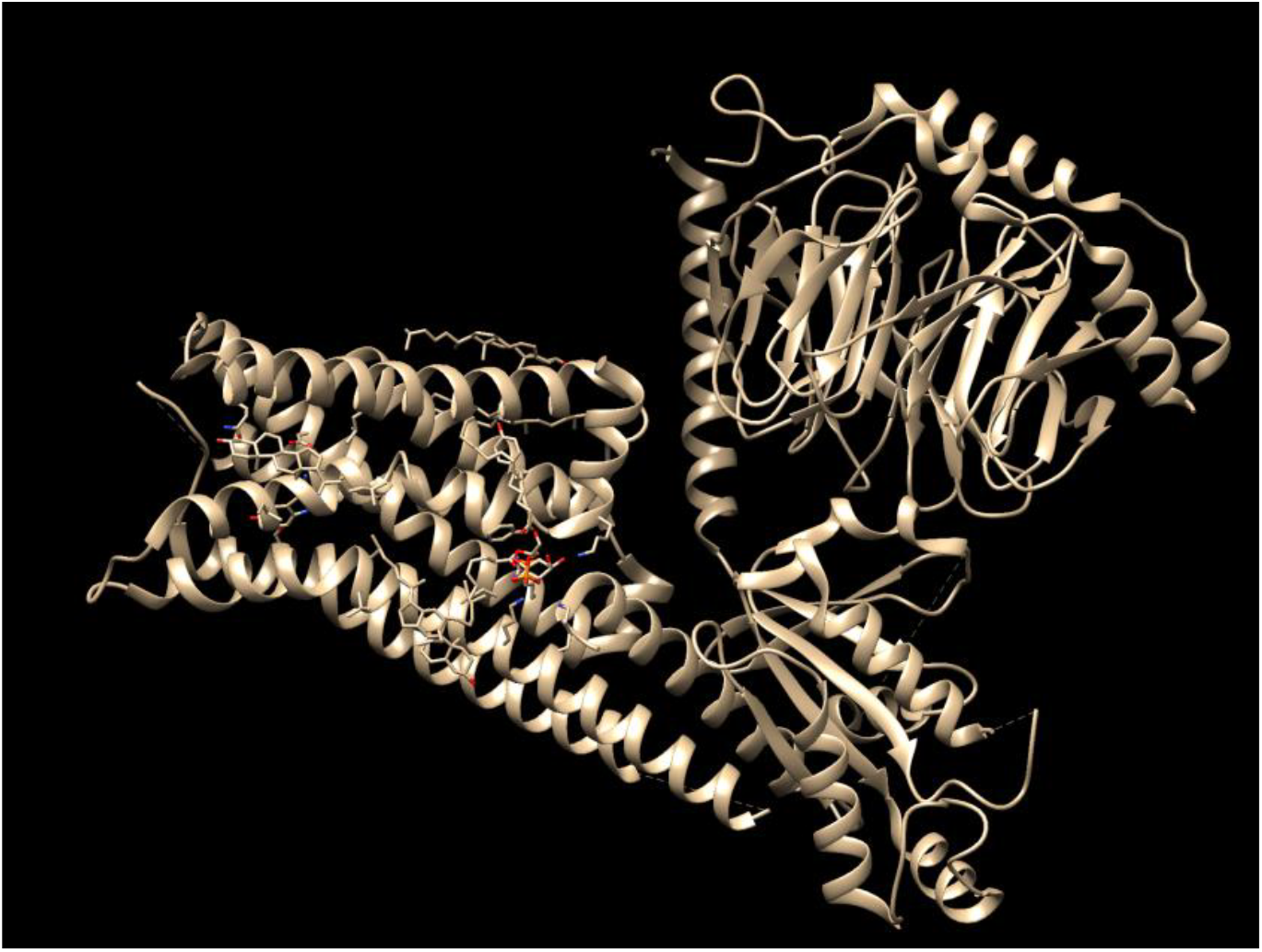
3D Structure of 5-HT1A (PDB:7E2Y). Image generated from Chimera.

Table 2 and Table 3 outline the binding affinity the ligands have to the protein of interest. Buspirone had the highest binding affinity to chain A, with a calculated value of -6.141 kcal/mol. However, as seen in chain G, there is very low binding affinity that can be deducted from the calculated affinity of 0 kcal/mol. Tandospirone had similar results, where chain A represented the highest affinity binding region at a calculated kcal/mol of -5.882. Chain B, yielded an affinity close to Chain A at -5.335 kcal/mol. However, just as Buspirone, chain G did not produce results that align with high affinity..

**Table 1:** Molecular docking data for ligand Buspirone generated from SwissDock.

| Buspirone Docking |  |  |  |  |  |  |  |
| --- | --- | --- | --- | --- | --- | --- | --- |
| HTR1A Chain A |  | HTR1A Chain B |  | HTR1A Chain G |  | HTR1A Chain R |  |
| Model | Calculated Affinity kcal/mol | Model | Calculated Affinity kcal/mol | Model | Calculated Affinity kcal/mol | Model | Calculated Affinity kcal/mol |
| 1 | -6.141 | 1 | -5.075 | 1 | 0 | 1 | -4.61 |
| 2 | -6.107 | 2 | -5.046 | 2 | 0 | 2 | -4.536 |
| 3 | -6.035 | 3 | -5.036 | 3 | 0 | 3 | -4.477 |
| 4 | -5.976 | 4 | -4.896 | 4 | 0.001 | 4 | -4.42 |
| 5 | -5.632 | 5 | -4.817 | 5 | 0.001 | 5 | -4.377 |
| 6 | -5.579 | 6 | -4.756 | 6 | 0.004 | 6 | -4.35 |
| 7 | -5.329 | 7 | -4.715 | 7 | 0.055 | 7 | -4.321 |
| 8 | -4.985 | 8 | -4.711 | 8 | 0.062 | 8 | -4.305 |
| 9 | -4.907 | 9 | -4.711 | 9 | 0.102 | 9 | -4.272 |
| 10 | -4.781 | 10 | -4.522 | 10 | 0.119 | 10 | -4.235 |
| 11 | -4.71 | 11 | -4.502 | 11 | 0.12 | 11 | -4.184 |
| 12 | -4.64 | 12 | -4.495 | 12 | 0.121 | 12 | -4.027 |
| 13 | -4.566 | 13 | -4.351 | 13 | 0.14 | 13 | -3.949 |
| 14 | -4.418 | 14 | -4.325 | 14 | 0.204 | 14 | -3.935 |
| 15 | -4.272 | 15 | -4.112 | 15 | 0.207 | 15 | -3.917 |
| 16 | -4.246 | 16 | -3.974 | 16 | 0.226 | 16 | -3.868 |
| 17 | -4.135 | 17 | -3.972 | 17 | 0.268 | 17 | -3.863 |
| 18 | -4.095 | 18 | -3.966 | 18 | 0.316 | 18 | -3.833 |
| 19 | -4.082 | 19 | -3.913 | 19 | 0.381 | 19 | -3.766 |
| 20 | -3.939 | 20 | -3.91 | 20 | 0.448 | 20 | -3.656 |

**Table 2:** Molecular Docking Data for ligand Tandospirone generated from SwissDock.

| Tandospirone Docking |  |  |  |  |  |  |  |
| --- | --- | --- | --- | --- | --- | --- | --- |
| HTR1A Chain A |  | HTR1A Chain B |  | HTR1A Chain G |  | HTR1A Chain R |  |
| Model | Calculated Affinity kcal/mol | Model | Calculated Affinity kcal/mol | Model | Calculated Affinity kcal/mol | Model | Calculated Affinity kcal/mol |
| 1 | -5.882 | 1 | -5.355 | 1 | 0 | 1 | -4.862 |
| 2 | -5.808 | 2 | -5.236 | 2 | 0 | 2 | -4.782 |
| 3 | -5.768 | 3 | -5.221 | 3 | 0.001 | 3 | -4.739 |
| 4 | -5.586 | 4 | -5.096 | 4 | 0.001 | 4 | -4.548 |
| 5 | -5.504 | 5 | -5.029 | 5 | 0.001 | 5 | -4.488 |
| 6 | -5.151 | 6 | -5.019 | 6 | 0.002 | 6 | -4.437 |
| 7 | -4.921 | 7 | -4.873 | 7 | 0.002 | 7 | -4.326 |
| 8 | -4.774 | 8 | -4.743 | 8 | 0.002 | 8 | -4.282 |
| 9 | -4.611 | 9 | -4.668 | 9 | 0.002 | 9 | -4.28 |
| 10 | -4.596 | 10 | -4.566 | 10 | 0.002 | 10 | -4.214 |
| 11 | -4.297 | 11 | -4.532 | 11 | 0.002 | 11 | -4.088 |
| 12 | -4.097 | 12 | -4.48 | 12 | 0.002 | 12 | -4.043 |
| 13 | -3.997 | 13 | -4.479 | 13 | 0.002 | 13 | -3.782 |
| 14 | -3.845 | 14 | -4.469 | 14 | 0.002 | 14 | -3.755 |
| 15 | -3.844 | 15 | -4.341 | 15 | 0.003 | 15 | -3.745 |
| 16 | -3.824 | 16 | -4.098 | 16 | 0.003 | 16 | -3.726 |
| 17 | -3.773 | 17 | -3.997 | 17 | 0.003 | 17 | -3.658 |
| 18 | -3.563 | 18 | -3.882 | 18 | 0.003 | 18 | -3.653 |
| 19 | -3.503 | 19 | -3.874 | 19 | 0.004 | 19 | -3.572 |
| 20 | -3.343 | 20 | -3.858 | 20 | 0.006 | 20 | -3.503 |

### Toxicity Prediction

The raw data from ProTox 3.0 predicted Buspirone would be in a toxicity class of 3 and the predicted LD50 is 196mg/kg. Figure 4 details the possible active toxic regions for this molecule. The highest probability (0.89) of toxicity is neurotoxicity, followed by respiratory, blood brain barrier, and clinical toxicity.

**Figure 4.**
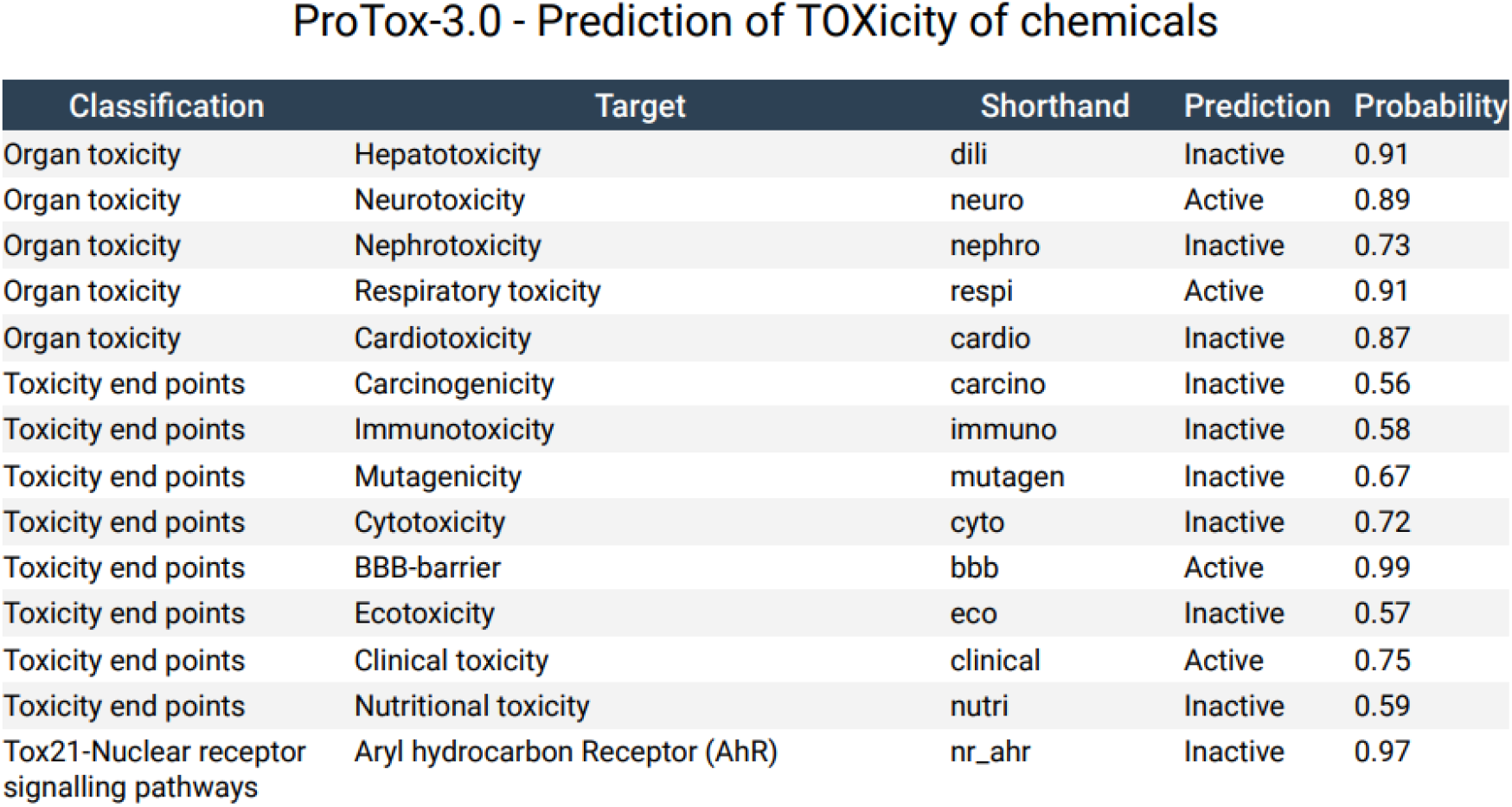
Predicted toxicity report for Buspirone from ProTox 3.0. Only two classifications are included in the figure.

Buspirone exhibits high probabilities for neurotoxicity (0.89) and respiratory toxicity (0.91), suggesting that these are the most significant concerns (Fig.4). The high probability for BBB penetration (0.99) indicates that the drug effectively crosses into the brain, which is necessary for its action but also raises concerns about CNS-related side effects (Fig.4). The clinical toxicity probability (0.75) indicates a moderate to high risk of adverse effects under therapeutic conditions (Fig.4).

Alternatively the raw data from ProTox 3.0 for Tandospirone predicted it to be in toxicity class 4 and have a LD50 of 590 mg/kg. Figure 5 depicts the probability of active target toxicity. The highest probability is for neurotoxicity followed by respiratory, immuno, blood brain barrier and clinical toxicity. Figure 6 visually demonstrated the dose value distributions for both compounds. Tandospirone shows high probabilities for neurotoxicity (0.89) and respiratory toxicity (0.89) (Fig.5). Notably, the BBB penetration probability is 1.00, confirming that Tandospirone also readily crosses into the CNS (Fig.5). Additionally, Tandospirone presents a significant probability for immunotoxicity (0.73), which suggests that immune system-related side effects should be considered during treatment (Fig.5). Similar risk profile of complications is indicated for Tandospirone, compared to Buspirone.

**Figure 5.**
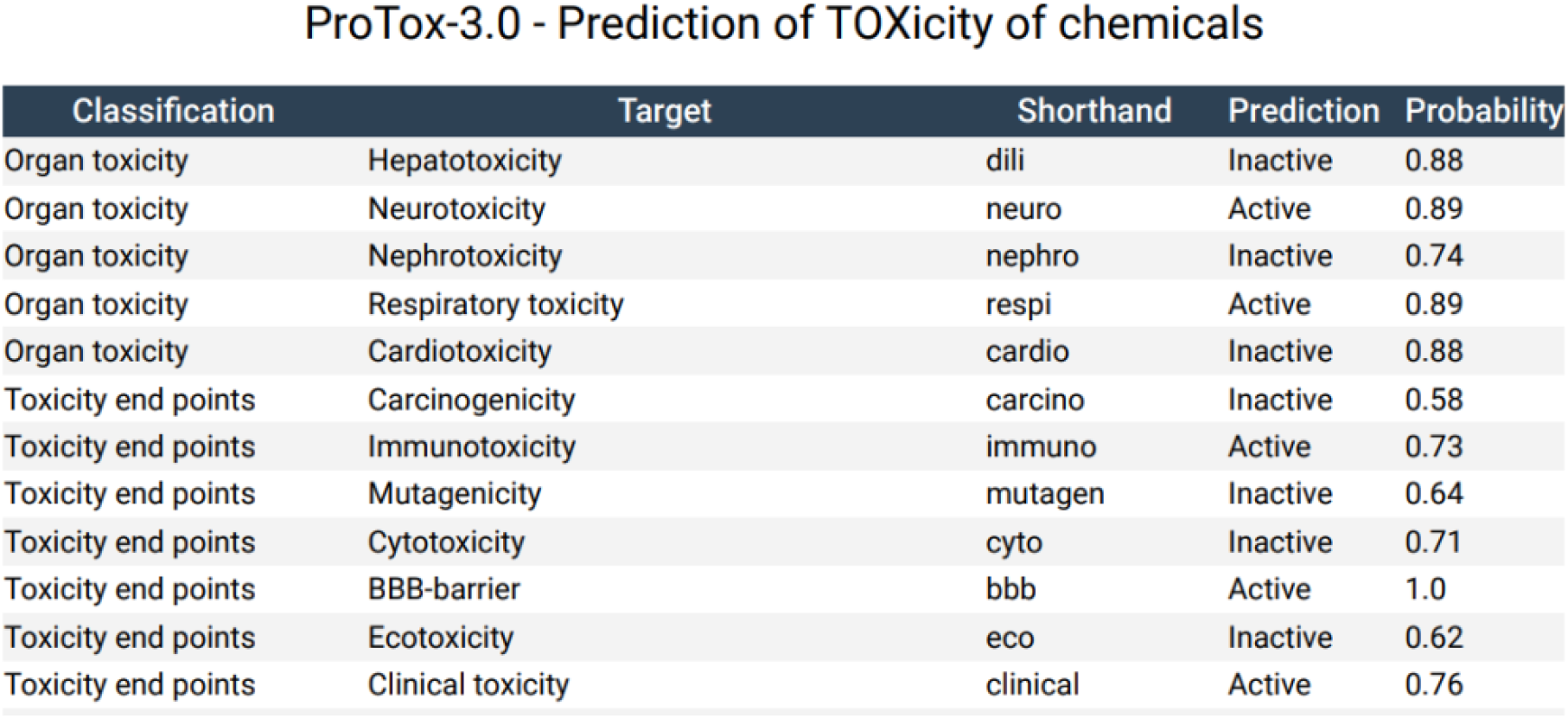
Predicted toxicity report for Tandospirone from ProTox 3.0. Only two classifications are included in the figure

**Figure 6.**
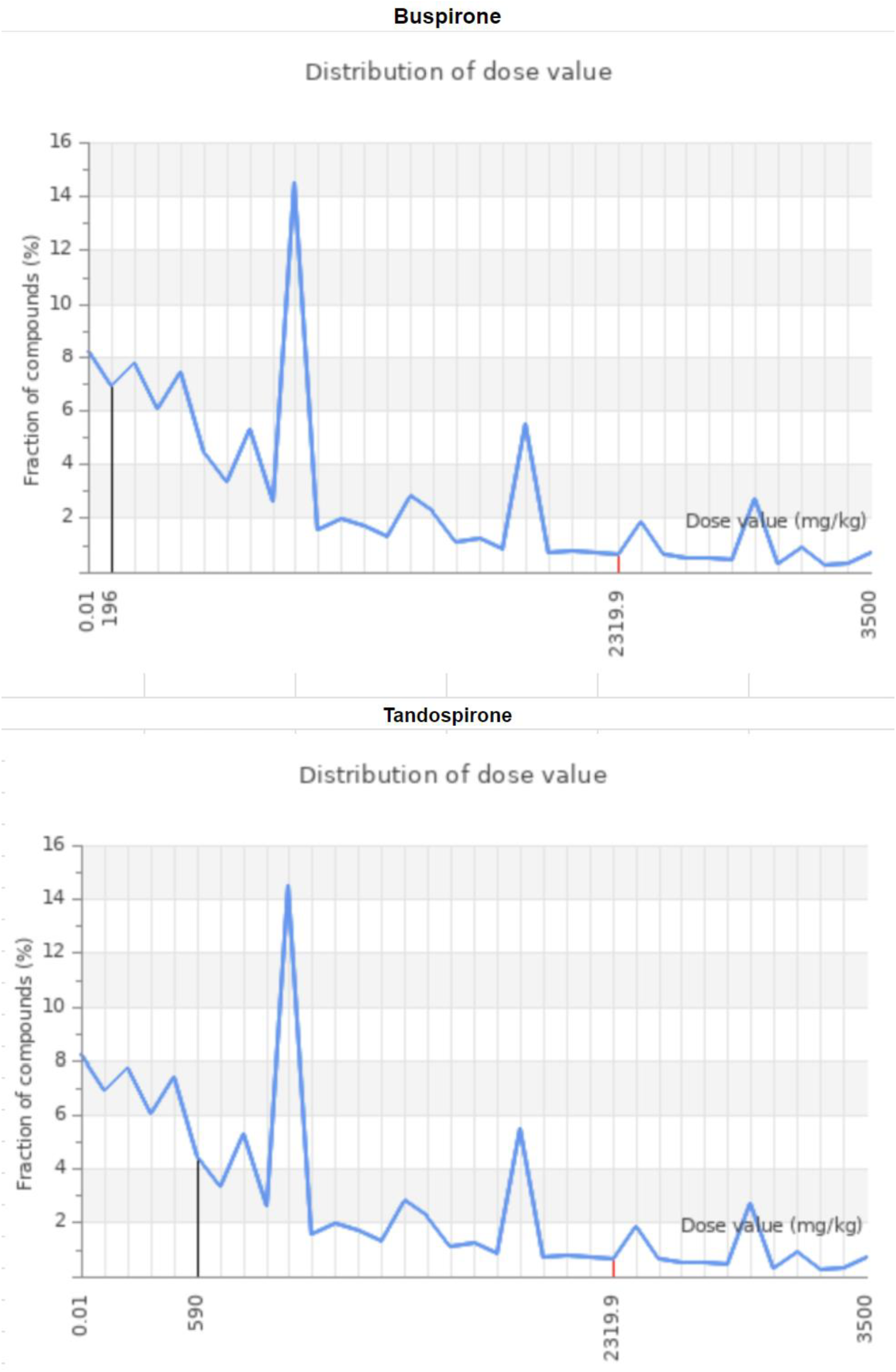
Distributions of dose values generated from ProTox for Buspirone and Tandospirone

## 6 DISCUSSION & INTERACTION ANALYSIS

### INTERACTION ANALYSIS

The molecular docking analysis of Buspirone and Tandospirone with the HTR1A receptor has indicated some interaction dynamics. Buspirone demonstrated the highest binding affinity with Chain A, with a calculated value of -6.141 kcal/mol. This strong and stable interaction suggests that Chain A represents a receptor conformation that is particularly favorable for Buspirone binding. Since Buspirone is a known serotonin receptor agonist, this interaction is crucial for its role in modulating serotonin levels and effectively treating anxiety disorders. The binding in Chain A may involve key residues that facilitate optimal hydrogen bonding and hydrophobic interactions. This high binding affinity ensures a stable ligand-receptor complex that enhances serotonin signaling (Jóźwiak & Płazińska., 2021). Conversely, the lower binding affinities were observed in Chains G and R for Buspirone. Some models, however, showed positive values, referring to the suboptimal or destabilizing interactions. These findings suggest that Buspirone may not effectively engage with the receptor in these conformations. It may be partially due to steric hindrances or the lack of critical contact points necessary for stable binding (Wang et al., 2013). These findings collectively indicate that different receptor chains exhibit different binding affinities within the HTR1A receptor and different conformations can significantly influence the drug’s efficacy, with Chain A holds the greatest potential for effective binding.

Similarly, Tandospirone exhibited the highest binding affinity with Chain A, calculated at -5.882 kcal/mol. Although slightly lower than Buspirone’s affinity in Chain A, this result indicates that Tandospirone also effectively interacts with the HTR1A receptor in this conformation. The interaction likely involves stabilizing forces similar to those observed with Buspirone, such as hydrogen bonds and hydrophobic contacts, which are essential for receptor activation. However, compared to Buspirone, the binding affinities of Tandospirone in Chains G and R were significantly lower, with positive values in some models. This suggests that Tandospirone’s efficacy is highly dependent on the receptor’s conformation, and the lower affinities in these chains may reflect conformational constraints or suboptimal binding environments, and they do not favor Tandospirone’s interaction.

The predicted toxicity analysis elucidated that there are potential risk factors associated with Busiprone and Tandospirone. However, those risk factors are at incredibly high doses of the drug. The average Busiprone dose for the adults with anxiety ranges from 7.5 mg to no more than 60 mg daily (Mayo Clinic 2024). PubChem reports that the compound has not been linked to significant toxicity. The effects of the drug can cause inconsistent serum aminotransferase elevation but there is no clinical evidence that the liver is damaged (PubChem 2024). Tandospirone however is still experimental but studies suggest the dose should not exceed 60 mg daily (Li et al., 2022). There is little evidence supporting the toxicity levels of Tandospirone but it has been indicated by PubChem that there is warning of target organ toxicity and respiratory irritation, which supports our prediction studies.

The predicted toxicity profiles for Buspirone and Tandospirone are essential when considering for treating anxiety and mood disorders. Both compounds have high probabilities of neurotoxicity (0.89) and respiratory toxicity (0.91 for Buspirone and 0.89 for Tandospirone), which align with their roles as HTR1A agonists primarily targeting the CNS. These findings are not surprising given the mechanism of action of these drugs, including the modulation of serotonin levels within critical brain regions responsible for mood regulation (Le François et al., 2008). However, the strong BBB penetration probabilities (0.99 for Buspirone and 1.00 for Tandospirone) warrant more careful inspection. While effective CNS penetration is necessary for these drugs to exert their therapeutic effects, it also raises concerns about the potential for CNS-related side effects. The high neurotoxicity that these drugs have been predicted pose a risk.It projected that patients could experience potential adverse neurological effects at higher doses or with prolonged use of the medication. The careful dosage management and monitoring strategy are crucial for optimal therapeutic effect (Chang et al., 2014).

Furthermore, Tandospirone’s significant immunotoxicity probability (0.73) indicates a potential impact on the immune system, which might manifest as immune-related adverse events. This is particularly relevant given the growing recognition of the interplay between the immune system and mental health, as immune dysregulation could worsen the psychiatric disorder(Ford & Savitz., 2023).By recognizing these risks early in the drug discovery process, researchers can modify the chemical structures of these ligands to reduce toxicity while maintaining efficacy with the consideration of the immune system involvement, leading to safer and more effective treatments.

Both Buspirone and Tandospirone show strong binding to Chain A of the HTR1A receptor, the lower affinities in Chains G and R, coupled with the toxicity risks. However, on a more positive note, the differential binding affinities and predicted toxicity profiles suggest that there is substantial room for drug optimization. For instance, modifying the molecular structures to enhance binding in less favorable receptor conformations could increase the overall therapeutic efficacy while potentially reducing off-target effects (Congreve et al., 2020).

## 7 CONCLUSION

In summary, the molecular docking studies of Buspirone and Tandospirone with the HTR1A receptor concluded the critical role of Chain A in facilitating effective binding, highlighting its therapeutic potential in modulating serotonin levels to treat anxiety and mood disorders. The observed variability in binding affinities across different receptor chains emphasize the importance of conformation in terms of drug efficacy. Given the potential risks of neurotoxicity, respiratory toxicity and BBB toxicity associated with these drugs predicted by the toxicity profile, the adverse side effect should also be an aspect worth careful examination. Structural modifications to enhance binding affinities in less favorable receptor conformations could improve the overall therapeutic efficacy while reducing off-target effects. Such advancements would contribute to the development of more precise and effective treatments for anxiety and mood disorders.

## DATA AVAILABILITY

**Chemical Entities of Biological Interest (ChEBI)**

(https://www.ebi.ac.uk/chebi/)

**Ensembl**

(https://useast.ensembl.org/index.html)

**NCBI (National Center for Biotechnology Information):**

(https://www.ncbi.nlm.nih.gov/)

**ProTox-3.0 Prediction of TOXicity in Chemicals**

(https://tox.charite.de/protox3/index.php?site=compound_input)

**SwissDock**

(https://www.swissdock.ch/)

**UCSF ChimeraX**

(https://www.cgl.ucsf.edu/chimerax/

